# SIRT6 activator fucoidan extends healthspan and lifespan in aged wild-type mice

**DOI:** 10.1101/2025.03.24.645072

**Authors:** Seyed A. Biashad, Eric Hillpot, Francesco Morandini, Cheyenne Rechsteiner, Victoria Paige, Gregory Tombline, Minseon Lee, Zhizhong Zheng, Yuan Liang, John Martinez, Natasha Sieczkiewicz, Zhihui Zhang, Valentin Volobaev, Denis Firsanov, Matthew Simon, Lei J. Zhang, Paul D. Robbins, Andrei Seluanov, Vera Gorbunova

**Author notes:** equal contribution (Seyed A. Biashad, Eric Hillpot, Francesco Morandini, Cheyenne Rechsteiner).

## Abstract

SIRT6 is a protein deacylase, deacetylase, and mono-ADP-ribosylase (mADPr) regulating biological pathways important for longevity including DNA repair and silencing of LINE1 retrotransposons. SIRT6 knockout mice die by 30 days of age, whereas SIRT6 overexpression increases lifespan in male mice. Finding safe pharmacological activators of SIRT6 would have clinical benefits. Fucoidan, a polysaccharide purified from brown seaweed, has been identified as an activator of SIRT6 deacetylation activity. Here, we show that fucoidan also activates SIRT6 mADPr activity, which was shown to be elevated in certain human centenarians. Administering fucoidan to aged mice led to a significant increase in median lifespan in male mice. Both male and female mice demonstrated a marked reduction in frailty and epigenetic age. Fucoidan-treated mice showed repression of LINE1 elements suggesting that the beneficial effects of fucoidan are mediated, at least in part, by SIRT6. As brown seaweed rich in fucoidan is a popular food item in South Korea and Japan, countries with the highest life expectancy, we propose that fucoidan supplementation should be explored as a safe strategy for activating SIRT6 and improving human healthspan and lifespan.

## Introduction

Sirtuins are NAD+-dependent deacetylases/deacylases and ADP-ribosyltransferases involved in the regulation of organismal lifespan in numerous species ranging from yeast to mammals^1,2^. SIRT6, a mammalian sirtuin, regulates gene expression, metabolism, epigenome maintenance, and genomic stability. Under conditions of cellular oxidative stress, SIRT6 is phosphorylated by JNK^3^ and promotes DNA repair by mono-ADP ribosylation and activation of PARP1^4^. Additionally, the ability of overexpressed SIRT6 to promote DNA double-strand break repair correlates with species lifespan in rodents^5^. SIRT6 knockout mice have a severely shortened lifespan characterized by genomic instability and defects in DNA repair, as well as broad degenerative phenotypes and metabolic defects^6^. Mice deficient in SIRT6 also experience de-repression of LINE1 transposons, which causes an inflammatory response through the accumulation of cytoplasmic LINE1 cDNA^7^. Conversely, mice overexpressing SIRT6 experience extended lifespan and reduced frailty^8^. As such, SIRT6 pharmacological activation is a promising therapeutic target to alleviate age-related frailty and extend lifespan.

SIRT6 has several related but distinct catalytic activities and targets. These include histone deacetylase activity on H3K9^9^, H3K18^10^, and H3K56^11,12^. Additionally, SIRT6 can act as a protein mono-ADP ribosyltransferase, with multiple known substrates involved in chromatin modification and DNA repair, including PARP1^4^, TRIM28 (KAP1)^13^, KDM2^14^, and BAF170^15^. Remarkably, a SIRT6 variant enriched in human centenarians was shown to have enhanced mono-ADP-ribosylation (mADPr) activity, pointing toward a specific importance of this activity in the longevity-promoting effects of SIRT6 ^16^.

Prior studies have identified several small molecule activators of SIRT6 deacetylation activity that showed beneficial biological effects in cell culture. Small-molecule allosteric activators of SIRT6 deacetylation MDL-800 and MDL-801 exerted antiproliferative effects in non-small cell lung carcinoma cells^17^ and improved genome stability^18^. The related activator MDL-811 alleviated neuroinflammation in mice^19^. A few natural polyphenolic compounds, including quercetin and cyanidin, have also been reported to activate SIRT6 deacetylation activity *in vitro*^20,21^. No chemical activators of SIRT6 mADPr activity have been reported.

Fucoidan is a complex sulfated polysaccharide found in brown algae and other marine organisms where it serves a structural and protective role. The main monomeric subunit of the fucoidan polymer is the monosaccharide fucose, although other monosaccharides are also incorporated into fucoidan in smaller quantities. Seaweed and fucoidan have a long history as traditional remedies and continue to be used as complementary medicine for various ailments today in countries such as Japan and South Korea. Fucoidan was found to have many therapeutic properties^22^. Fucoidan is efficiently absorbed through the intestine in humans^23^ and oral supplementation of fucoidan increases muscle size and strength^24,25^, mitigates liver injury^26,27^, enhances immunity and alleviates inflammation^28^ in mouse models. Fucoidan also showed health benefits in humans including improved quality of life in cancer patients and improved response to influenza vaccine in elderly people^29–32^. Fucoidan was previously identified as an activator of SIRT6 deacetylase activity^29^, but its effect on the mADPr activity of SIRT6 has not been examined. It is plausible that the reported beneficial effects of fucoidan may be mediated by activation of SIRT6 and mitigation of inflammation.

Here, we report that fucoidan extract from *Fucus vesiculosus* strongly stimulates the SIRT6 mADPr activity. Furthermore, fucoidan dietary supplementation extends the lifespan of C57BL/6 male mice and promotes health in both sexes. At the cellular and molecular levels, we observe silencing of LINE1 elements, reduced inflammation, induction of pathways involved in DNA repair and chromatin organization, and improved stress resistance.

## Results

### Fucoidan stimulates SIRT6 mADPr activity

As SIRT6 mADPr activity is specifically enhanced in certain human centenarians with rare coding variants in SIRT6^16^, we sought to identify activators of this activity. We tested several molecules previously reported to stimulate SIRT6 deacetylase activity using our mADPr assay^4^. The tested compounds were added to *in vitro* reactions containing purified SIRT6 and ^32^P-labeled NAD+, and SIRT6 mADPr activity was quantified by autoradiography. Of the compounds tested, only fucoidan increased SIRT6 mADPr activity (t-test: p = 0.005, **Figure 1A**). Interestingly, most of the other SIRT6 deacetylase activators showed an inhibitory effect. We also tested the effect of fucoidan on the ability of SIRT6 to deacetylate H3K9ac in nucleosomes, and found that fucoidan stimulated SIRT6 deacetylase activity, as reported previously^29^ (**Figure 1B**). We next compared several different fucoidan extracts from different seaweed species and sources to determine their relative effects on SIRT6 mADPr activity. Highly purified extract of *F. vesiculosus* showed the strongest stimulation of ribosylation (t-test: p = 0.001) with highly purified extracts from *U. pinnatifida* and *C. okamuranus* as well as nutrition supplement grade *F. vesiculosus* fucoidan supplied by DoNotAge (D.N.A.) also showing stimulation of ribosylation activity (t-test: p = 0.016, p = 0.016 and p = 0.028, **Figure 1C**). Titration of fucoidan extract concentrations showed that stimulation of mADPr activity approached saturation at concentrations of 0.2-0.5 mg/mL (**Figure 1D**). L-fucose, the sugar monomer found in fucoidan, had very mild stimulatory effect on SIRT6 deacetylation activity and no significant effect on mADPr activity (**Figure 1B, C**).

**Figure 1:**
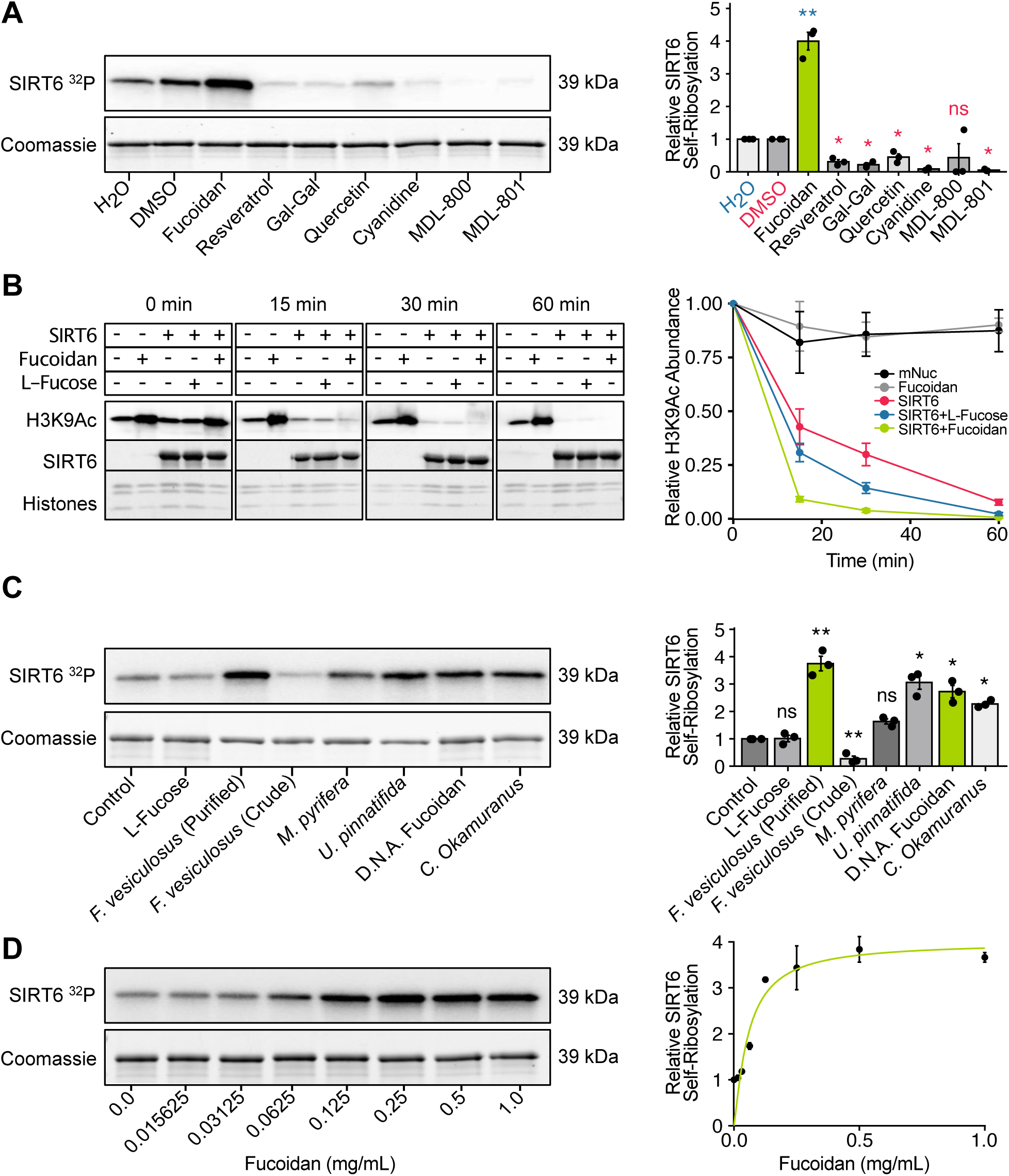
Fucoidan increases Sirt6 deacetylase and mADPr activity. **(A)** Test of Sirt6 self-mADPr activity by incorporation of ^32^P labeled NAD+, in reactions with various candidate activators. Representative gel is shown on the left, and quantification of three independent experiments is shown on the right. **(B)** Effect of fucoidan and L-fucose on Sirt6 deacetylation kinetics. Representative gel is shown on the left, and quantification of four independent experiments is shown on the right. **(C)** Effect of various fucoidan extracts on Sirt6 self-mADPr activity measured by incorporation of ^32^P labeled NAD+. Representative gel is shown on the left, and quantification of three independent experiments is shown on the right. **(D)** Titration curve showing the effect of different fucoidan concentrations on Sirt6 mADPr activity. Representative gel is shown on the left, and quantification of three independent experiments is shown on the right. Stars indicate significance according to t-test (*** = p < 0.001, ** = p < 0.01, * = p < 0.05, ns = p ≥ 0.05), error bars represent standard errors of the mean.

In summary, most compounds that stimulate SIRT6 deacetylation had no or even inhibitory effects on mADPr, suggesting that these catalytic activities generally require different chemical activators. Fucoidan was the only tested compound displaying strong stimulatory effect of both deacetylation and mADPr activities of SIRT6.

### Fucoidan reduces frailty in aged mice and extends median lifespan of male mice

To assess the effects of fucoidan on the aging process *in vivo*, we administered fucoidan in drinking water and chow (a total average dose of 278 mg/day) to a cohort of 44 mice (22 male and 22 female) and 36 control mice starting at 15 months of age (**Figure 2A**). The mice were monitored for the remainder of their life to assess effects of fucoidan on lifespan. Additionally, mouse health was assessed using the Howlett frailty index^33^, between 19 and 25 months of age. Blood samples were collected from live animals at the ages of 22 and 26 months. For analysis of tissues, smaller cohorts of 5 male and 5 female mice were treated with fucoidan for 1 month before tissues were harvested.

**Figure 2:**
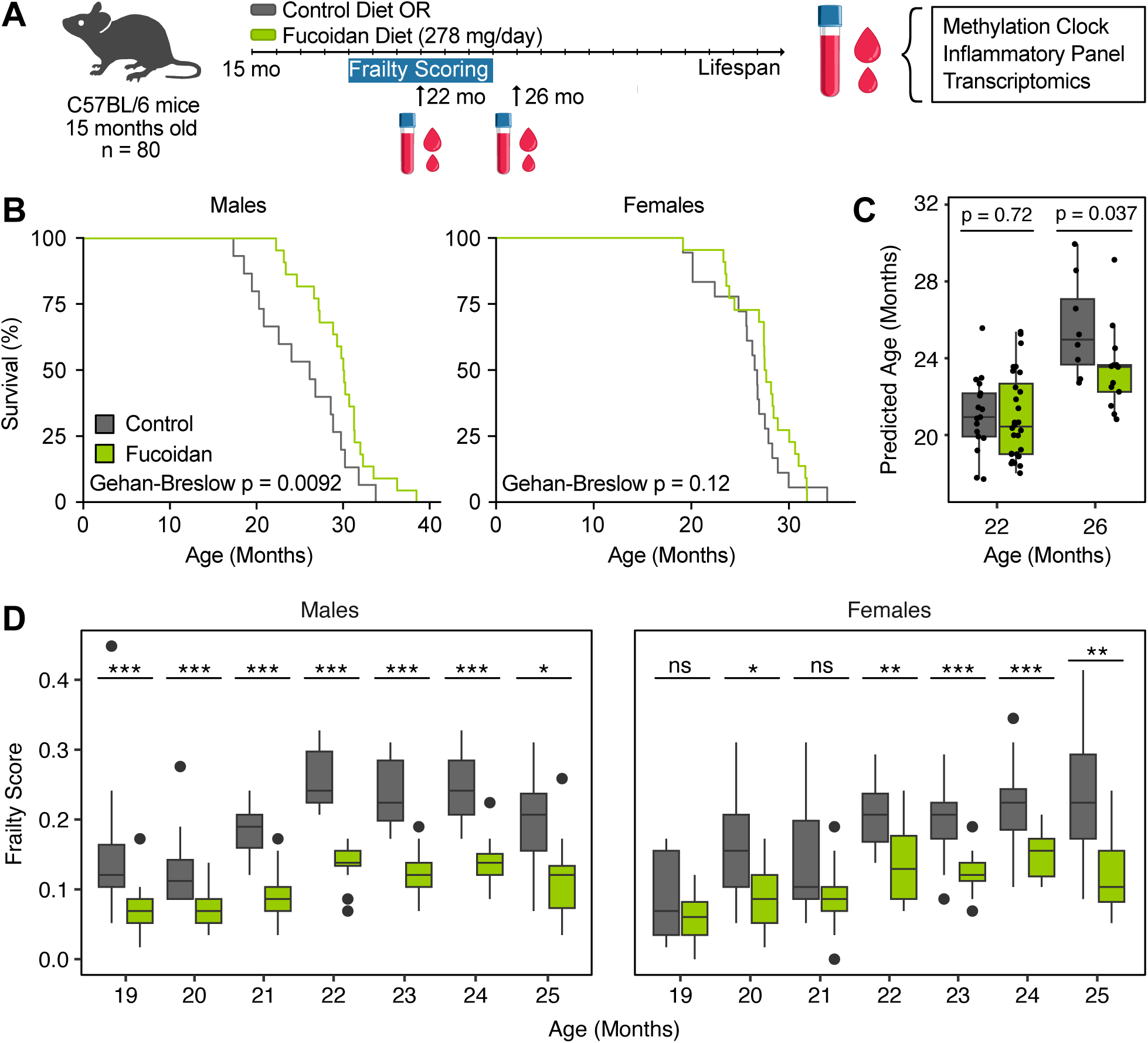
Fucoidan reduces frailty in aged mice and extends the lifespan of male mice. **(A)** Timeline of the survival study, frailty measurements and blood sample collection. **(B)** Survival curves for male and female mice fed with either control or fucoidan diets. P values were calculated with the Gehan-Breslow test. **(C)** Blood methylation age according to the UniversalClock3Blood clock. P-values were calculated with the Mann-Whitney U test. **(D)** Frailty scores of control and fucoidan fed mice. Stars indicate significance according to Mann-Whitney U tests within the time point (*** = p < 0.001, ** = p < 0.01, * = p < 0.05, ns = p ≥ 0.05). Overall significance: p = 0.046 for males, p = 0.017 for females (linear mixed model). The number of mice measured at each timepoint is shown in **figure S1B.**

Fucoidan treatment significantly extended the median lifespan of males by 13% (Gehan-Breslow test: p = 0.009, **Figure 2B**). Female lifespan, on the other hand, only showed a non-significant increase by 3% (Gehan-Breslow test: p = 0.12). Furthermore, fucoidan-treated mice of both sexes had lower DNA methylation age relative to controls at 26 months (Mann-Whitney U test, p = 0.037, **Figure 2C**). Fucoidan treatment did not have any effect on body mass (**Figure S1A**) indicating that mice did not change food intake in response to fucoidan in chow. This excludes health and longevity effects due to calorie restriction. Frailty scores for both males and females, assessed monthly starting at 19 months of age, increased at a slower rate in fucoidan-treated animals relative to controls (Linear mixed model, interaction term between diet and treatment time, p = 0.046 for males, p = 0.006 for females, **Figure 2D**). A breakdown of frailty components by organ/system revealed reduced frailty across nearly all systems in fucoidan-treated mice (**Figure S1B**).

To determine whether the lifespan-extending effects of fucoidan are SIRT6-mediated, we measured lifespan in SIRT6 knockout (SIRT6 KO) mice with or without fucoidan supplementation. In support of a SIRT6-mediated lifespan effect, no significant lifespan difference between fucoidan-fed and control SIRT6 KO mice could be observed (Gehan-Breslow test: p = 0.87, **Figure S2A**). We additionally looked for molecular evidence of increased SIRT6 activity *in vivo*. Detection of mADPr *in vivo* is complicated due to the absence of sensitive antibodies. Therefore, we measured acetylation of the SIRT6 substrate H3K9 (H3K9ac) in the lung and liver of mice treated with fucoidan for 1 month. Fucoidan treatment resulted in reduction of H3K9ac in the lungs of male mice (Mann-Whitney U test, p = 0.015, **Figure S2B**). Additionally, both lungs and liver of female mice showed a downward trend in H3K9ac (Mann-Whitney U test, p = 0.09, 0.072 respectively, **Figure S2B**).

In conclusion, fucoidan treatment provides systemic health benefits in both sexes, and extends the lifespan of male mice by 13%. These effects are likely to be mediated, at least in part, by enzymatic activation of SIRT6.

### Fucoidan downregulates aging and inflammation-related pathways and increases pathways related to DNA repair and chromatin maintenance

We performed RNA-seq on blood samples from 22 months old mice involved in the aging study. In males, 926 genes were upregulated, and 2272 genes were downregulated in fucoidan-treated versus control mice (**Figure 3A**), whereas in females, no gene reached the threshold for differential expression (**Figure S3A**). GSEA on Gene Ontology biological process terms revealed upregulation of terms related to RNA modification and B and T cell differentiation in males (**Figure 3B**). Downregulated terms were related to blood coagulation, myeloid identity, translation, stress and interferon response. Interestingly, fucoidan has been previously investigated as an anticoagulant, with promising results *in vivo* and *in vitro*^36–38^. ORA yielded similar results as GSEA, confirming upregulation of RNA processing pathways, T cell differentiation, and down regulation of coagulation and myeloid differentiation (**Figure 3C**). Additionally, ORA showed that upregulated genes were enriched for pathways that are regulated by SIRT6 such as DNA repair, telomere maintenance, histone modification and chromatin organization. Finally, the term “Aging” was downregulated in male mice, confirming the anti-aging effect according to the methylation clock. Coagulation was also downregulated in females, according to GSEA, however inflammatory pathways (response to lipopolysaccharide, IL-1ß and TNFα) were upregulated (**Figure S3B**).

**Figure 3:**
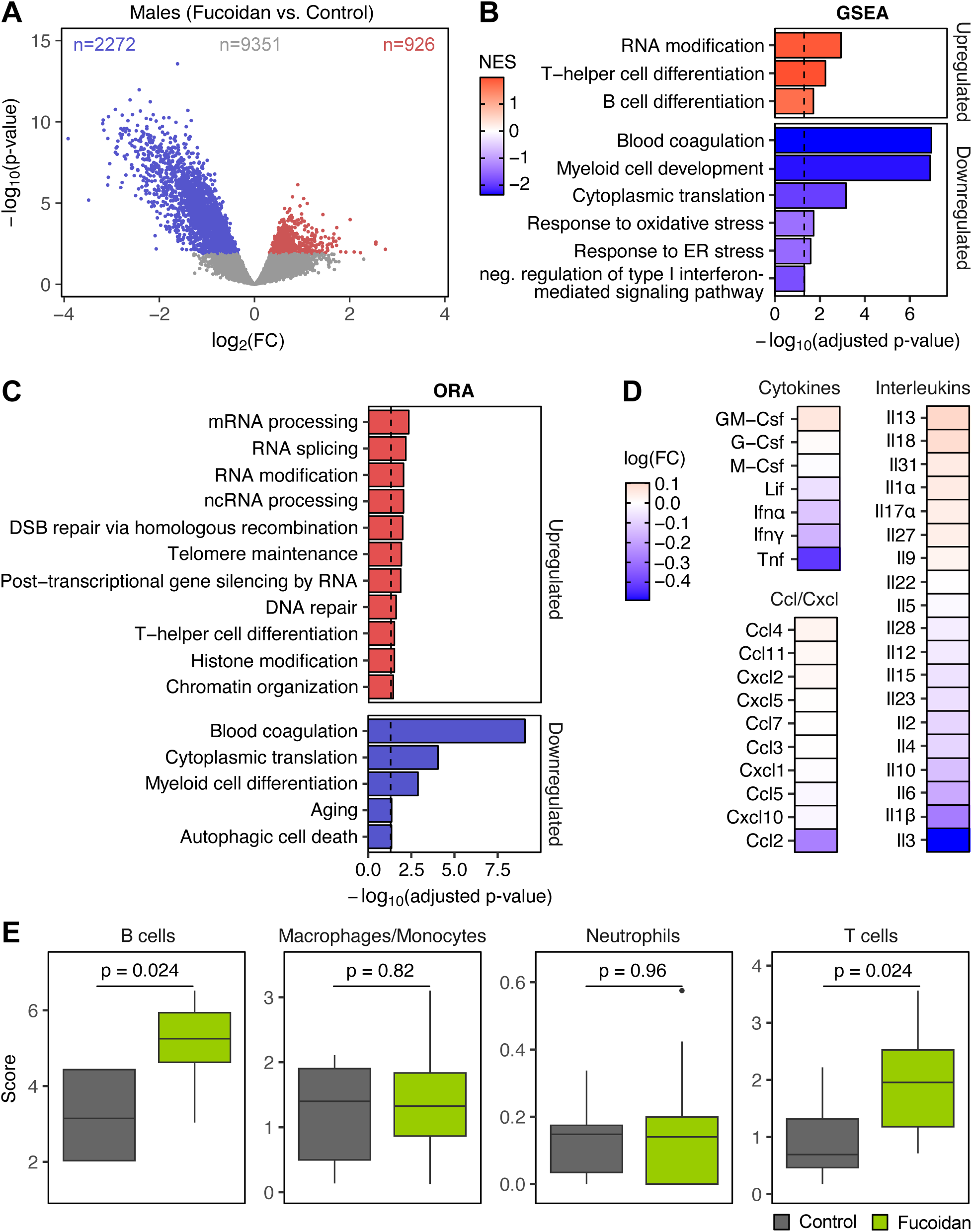
Fucoidan has an anti-aging and anti-inflammatory effect on the blood of male mice. **(A)** Volcano plot for the effect of fucoidan on 22-month male mice, which were part of the survival study. Colored dots denote significant up or downregulated genes (adjusted p-value < 0.05). **(B)** GSEA results showing GO terms significantly up or downregulated by fucoidan treatment in the blood of male mice. Selected terms are shown. **(C)** ORA results showing GO terms significantly enriched among up or downregulated genes in male mice. Selected terms are shown. **(D)** Plasma levels of cytokines in male mice measured by ELISA. **(E)** Estimation of blood cell composition in male mice by deconvolution using mMCP-counter. p-values were calculated with a Mann-Whitney U test.

As various terms related to lymphoid and myeloid cell identity were respectively upregulated and downregulated in males (**Figure 3B**), we asked if enrichment of these terms may be due to compositional changes in the blood rather than cell intrinsic effects. We used mMCP-counter^34^ to estimate blood cell composition from bulk data and found that male mice had a higher B cell and T cell fraction (Mann-Whitney U test, B cells: p = 0.024, T cells: p = 0.024, **Figure 3E**), whereas no major immune cell populations were different in females (**Figure S3D**). Fucoidan was previously shown to induce B and T cell expansion *in vitro*^35^. Aging is known to induce a myeloid bias in hematopoietic stem cell differentiation, which in turn contributes to impaired immune memory and sterile inflammation in aged individuals. Thus, the higher proportion of lymphoid cells in fucoidan-treated male mice reflect the anti-aging effect of fucoidan.

As terms related to interferon response (IFN) were downregulated in males (**Figure 3B**), we measured cytokines levels in the plasma using an ELISA panel. TNFα and IFNα and γ as well as Ccl2 and several pro-inflammatory interleukins such as IL-3, IL-1β, IL-6 tended to show lower abundance in the plasma of male fucoidan-treated compared to control mice (**Figure 3D**). Conversely, cytokine levels tended to be increased in female fucoidan-treated compared to control mice (**Figure S3C**), suggesting a sex-dependent effect of fucoidan on inflammatory profiles.

In summary, long-term treatment with fucoidan induced strong changes in the blood of male mice, including upregulation of SIRT-6 related pathways as DNA repair and chromatin maintenance and the associated downregulation of proinflammatory cytokines. Conversely, the blood of female mice showed much weaker differences, indicative of a mild inflammatory response. This discrepancy gives a possible explanation to the much larger lifespan extension in male animals.

### Fucoidan and Sirt6 overexpression exert a similar anti-inflammatory effect on male liver

Next, we performed RNA-seq on liver and lung samples collected after 1 month of fucoidan treatment starting at 15 months of age (**Figure 4A**). GO biological process analysis was then used on genes that were differentially expressed in the same direction across tissues or sexes (e.g., p.adj < 0.2 and logFC > 0, **Figure S4A, B**). Briefly, pathways upregulated in both male and female lungs were related to protein folding, while downregulated ones were related to coagulation, consistently with the results seen in blood. There was minimal overlap between genes upregulated in male and female livers, as well as between those downregulated in both. Given the liver’s central role in processing dietary metabolites, the differing responses to fucoidan treatment in male and female livers may explain the sexually dimorphic effects of fucoidan, including its impact on lifespan extension.

**Figure 4:**
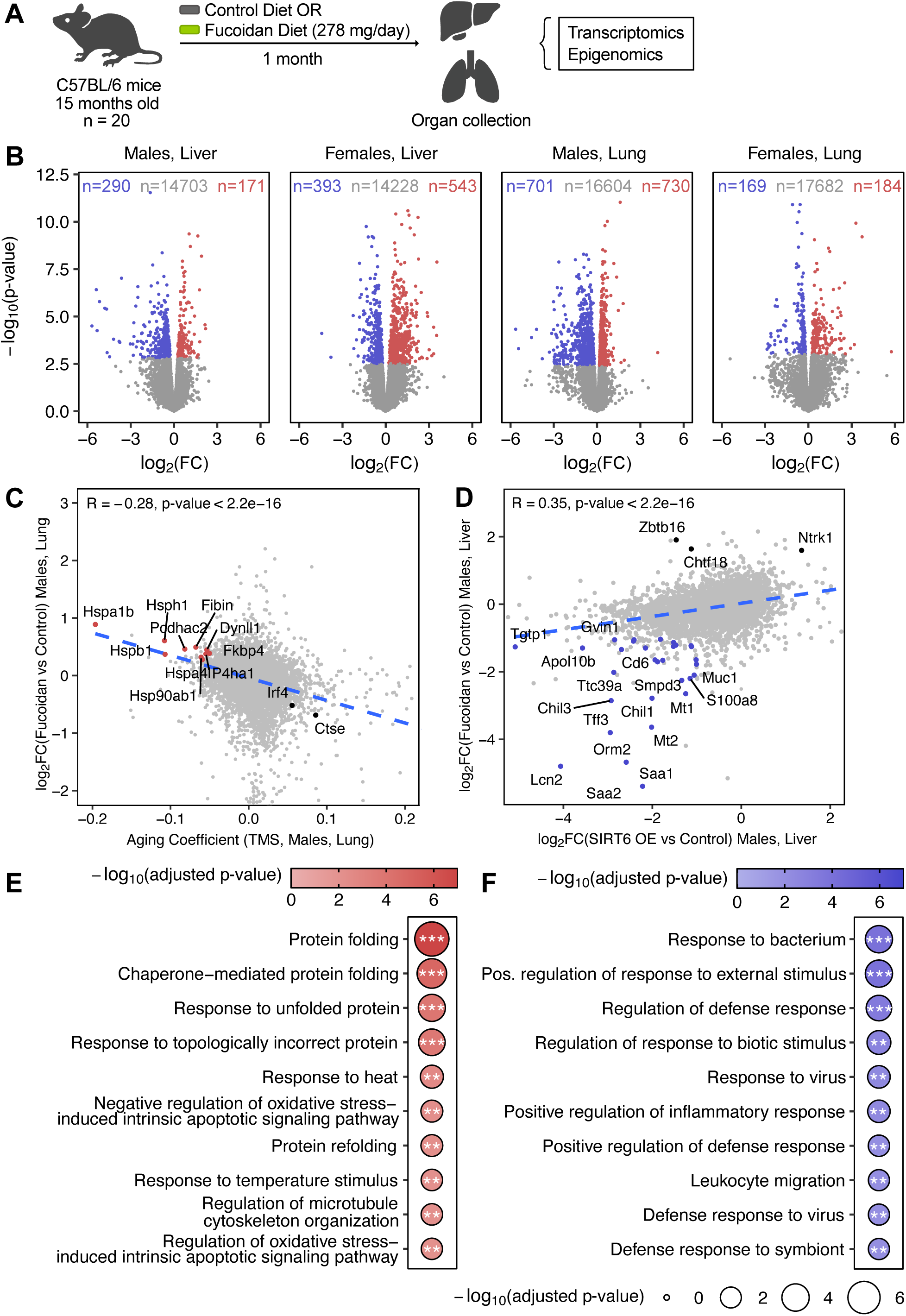
Comparison of the effects of fucoidan with those of aging and Sirt6 overexpression. **(A)** Experimental setup. Mice were treated with fucoidan or control diets starting at 15 month of age and liver and lung samples were collected after a month. **(B)** RNA-seq volcano plots for the effect of fucoidan on male and female liver and lung. Colored dots denote significant up or downregulated genes (adjusted p-value < 0.05). **(C)** Comparison of Fucoidan vs control logFCs with aging coefficients obtained from male lung samples from the Tabula Muris Senis bulk dataset. Darker dots represent genes differentially expressed in both Fucoidan vs control and during aging (adjusted p-value < 0.05 for aging, adjusted p-value < 0.1 for Fucoidan vs control). Red colored dots are genes significantly downregulated during aging and upregulated by Fucoidan. **(D)** Comparison of Fucoidan vs control logFCs with Sirt6 OE vs control logFCs obtained from liver data by Roichman *et al*. Darker dots represent genes differentially expressed in both Fucoidan vs control and Sirt6 OE vs control (adjusted p-value < 0.05, |logFC| > 1 for both). Blue colored dots are genes significantly downregulated in both experiments. **(E)** Gene ontology ORA for the genes highlighted in red in panel C. **(F)** Gene ontology ORA for the genes highlighted in blue in panel D.

To better understand the effects of fucoidan in the context of aging, we compared the changes in the tissues of fucoidan fed mice to bulk RNA-seq data from Tabula Muris Senis (TMS)^39^. We observed a negative correlation between the effects of aging in male lungs and those of fucoidan treatment (**Figure 4C, E**). Interestingly, multiple genes related to protein folding were downregulated during aging but upregulated by fucoidan treatment. The effect of fucoidan on male liver was not correlated with age-related changes (**Figure S5A**). Conversely, the effect of fucoidan on female liver and lung was positively correlated with the effect of aging (**Figure S5B, C**). Thus, it seems possible that fucoidan treatment may constitute a mild stress in females, at least with short-term treatment.

Finally, we compared the effect of fucoidan with that of Sirt6 overexpression, using liver Sirt6 overexpression data generated by Roichman *et al*^8^. In male liver, the effect of fucoidan treatment was positively correlated with that of Sirt6 overexpression, with both treatments inducing downregulation of pro-inflammatory genes and pathways (**Figure 4D, F**). Among the highly downregulated genes, Saa1/2 and Orm2 are part of the acute-phase response, which stimulates the liver to secrete various inflammatory and coagulant proteins in response to infection or injury. The acute phase response is primarily induced by circulating IL-6 and TNFα, which were reduced in our plasma data (**Figure 3C**). Once again, female liver showed the opposite, albeit weaker, trend (**Figure S5D**).

In conclusion, short-term treatment with fucoidan produced highly sexually dimorphic responses. Nonetheless, fucoidan upregulated protein folding and downregulated coagulation related pathways in both male and female lungs. Protein folding pathways were downregulated during aging in lung data from TMS, thus fucoidan appears to restore protein folding to more youthful levels. Finally, fucoidan and Sirt6 over-expression exerted a similar effect on male liver, resulting in the downregulation of inflammatory pathways, including the acute phase response.

### Fucoidan induces LINE1 silencing in lungs

SIRT6 plays important role in silencing LINE1 transposable elements by recruiting heterochromatin factors and promoting formation of repressive heterochromatin^13^. Sirt6 KO mice and cells exhibit LINE1 derepression, expression, and cytosolic LINE1 cDNA accumulation, resulting in chronic inflammation driven by the cGAS/STING cytosolic DNA sensor^40^. Similarly, LINE1 transposons undergo derepression during aging^40–42^ and silencing them has the potential to increase healthspan and lifespan.

Given the function of SIRT6 in silencing LINE1 elements, we quantified the expression of LINE1 families in our liver and lung data. Additionally, we performed ATAC-seq and MeDIP-seq to measure the chromatin accessibility and DNA methylation at LINE1s, respectively. In general, some LINE1s families were downregulated, and others were upregulated, following fucoidan treatment. However, in male and female lungs, expression of the vast majority of LINE1 families was downregulated, as highlighted by the asymmetry of the volcano plots (**Figure 5A**). The number of downregulated families was larger than what would be expected assuming equal likelihood of down- or up-regulation (Binomial test). Consistently, the majority of LINE1 families had reduced accessibility and increased DNA methylation in the lungs of fucoidan treated mice. The same analysis performed in liver tissue did not yield conclusive results (**Figure S5E**). In particular, some omic modalities indicated repression while others indicated derepression. This discrepancy may be due to different biases in terms of interrogated genomic regions between the techniques.

**Figure 5:**
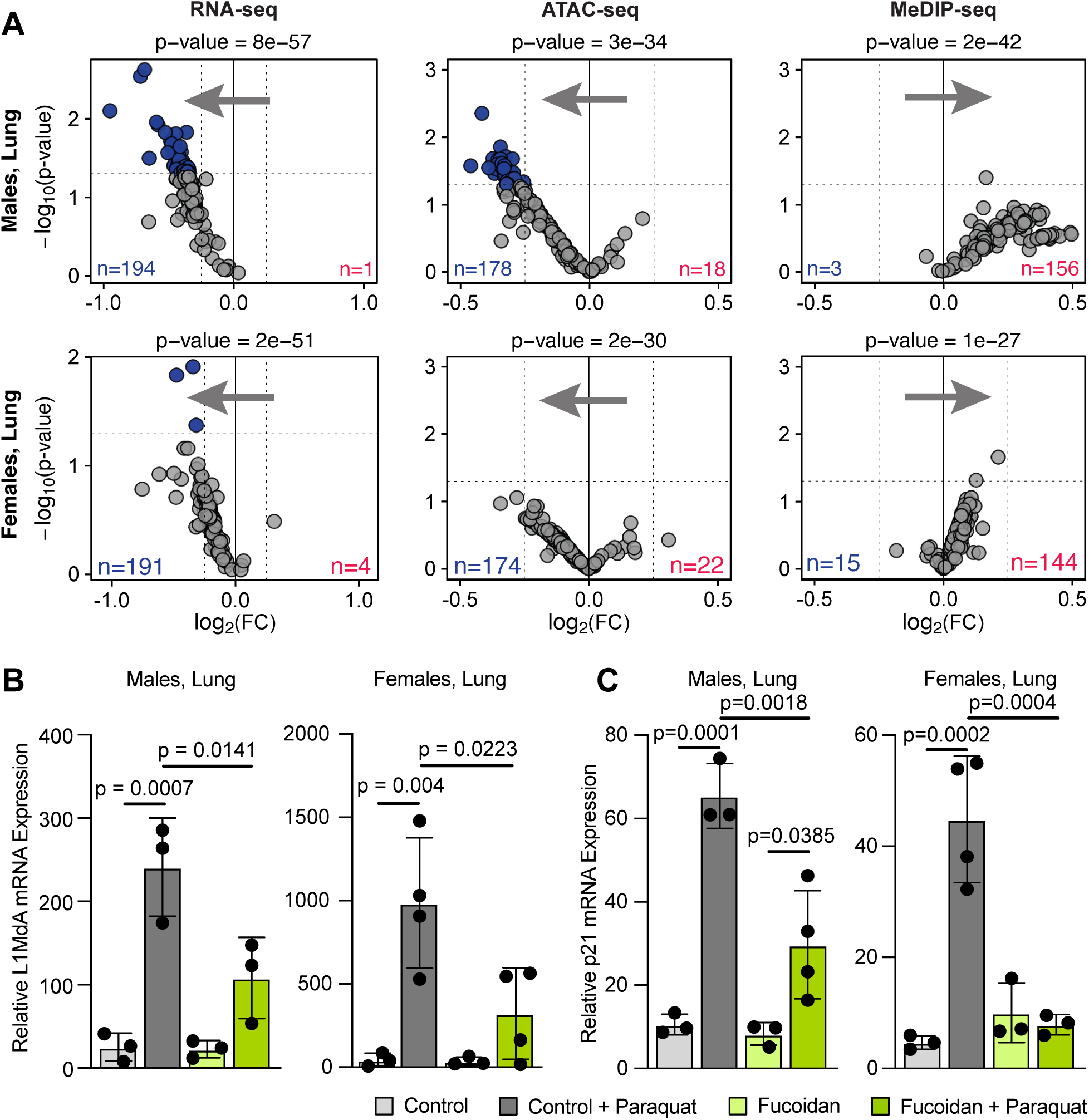
Fucoidan treatment induces LINE1 silencing in lungs basally and upon oxidative stress. **(A)** RNA-seq, ATAC-seq and MeDIP-seq volcano plots for the effect of fucoidan treatment on LINE1 element expression, accessibility and methylation in lung tissue. TE families with logFC < −0.25 and p < 0.05 (unadjusted) are colored in blue. TE families with logFC > 0.25 and p < 0.05 (unadjusted) are colored in red. Overall asymmetry of the volcano plot is tested for significant deviations from equal likelihood (50%) using a binomial test. **(B, C)** Quantification of LINE1 and p21 (CDKN1A) expression in lungs by RT-qPCR following oxidative stress by IP paraquat injection, in control mice or mice fed fucoidan diet for one month (3-4 biological replicates per group). P values were calculated using Tukey’s tests.

Transposable elements are known to undergo activation upon genotoxic stress^43^. We therefore tested whether fucoidan treatment ameliorates stress-induced LINE1 activation. We administered paraquat, an inducer of oxidative stress, to mice fed with fucoidan or control diet for 1 month. Paraquat caused strong upregulation of expression of the active LINE1 subfamily L1MdA in the lungs of control mice. This activation was significantly ameliorated in mice receiving fucoidan (**Figure 5B**). Interestingly, fucoidan-fed mice also showed lower levels of expression of *CDKN1a*, encoding the cyclin dependent kinase inhibitor p21^Cip1^, upon paraquat treatment (**Figure 5C**), suggesting that fucoidan has protective effect against genotoxic stress.

In summary, these results indicate that fucoidan treatment promotes silencing of active LINE1 elements in the lungs, resulting in reduced p21^Cip1^ levels, a marker of genotoxic stress. As SIRT6 promotes LINE1 silencing and DNA repair these results are consistent with the effect of fucoidan being mediated by SIRT6 activation.

## Discussion

As SIRT6 has been referred to as “the real aging Sirtuin”^44^, significant effort has been put into identifying chemical SIRT6 activators. Some of these molecules were shown to have beneficial effects in cell culture and *in vivo*^17–19^. Here we report that the SIRT6 activator, fucoidan, extends healthspan and lifespan *in vivo* in wild type mice, in particular, in male mice. Importantly, the extension was observed after starting fucoidan supplementation in middle-aged mice, suggesting that there is a benefit of initiating fucoidan treatment later in life. However, although there was a life extension only in male mice, fucoidan treatment led to a marked reduction in frailty in both male and female mice.

Other SIRT6 activators were identified based on their enhancing effect on deacetylase activity. In contrast, fucoidan activates both deacetylation and mADPr activities. Importantly, mADPr activity of SIRT6 is required for LINE1 inhibition^7^, a function which may be the key to reversing age-related inflammation and frailty. Consistently, we observed decreased inflammation in multiple tissues, and decreased LINE1 expression in lung tissue.

We show that not every species of brown algae produces fucoidan that activates SIRT6. Fucoidan is not a defined chemical and may consist of different ratios of fucose and galactose molecules which may be sulfated on different residues and form branched structures^45^. Hence only some species of brown algae produce fucoidan that has the chemical structure required for SIRT6 activation. As fucoidan is broken down in digestive tract, it is likely that a smaller, but not monomeric, fucoidan derivative with a defined chemical structure is responsible for SIRT6 activation. In the future, through carbohydrate chemistry, it should be possible to identify the minimal fucoidan unit with SIRT6 activating properties.

Fucoidan is remarkable because of its safety and the geographical locations where it is a food staple. Brown algae are a popular food in South Korea and Japan, the two countries with the longest life expectancies. Thus, fucoidan supplementation is a safe human intervention that may potentially improve quality of life in the elderly population. While a number of clinical trials had been conducted using fucoidan supplementation none of them had frailty or aging biomarkers as primary endpoints. Thus, a human clinical trial with fucoidan focused on healthspan and biomarkers of aging including analysis of LINE-1 expression and the epigenetic clock is needed.

### Limitations of the study

As fucoidan is a natural product with complex structure we cannot rule out that it has other cellular targets beyond SIRT6. However, as we show that fucoidan treatment affects SIRT6 targets such as H3K9ac and LINE1 transposons we conclude that at least some of the beneficial effects of fucoidan are mediated by SIRT6. Additionally, fucoidan exerted no beneficial effects on SIRT6 KO mice.

## Methods

### RESOURCE AVAILABILITY

#### Lead Contact

Further information and requests for resources and reagents should be directed to and will be fulfilled by the Lead Contact, Vera Gorbunova.

#### Materials Availability

All unique reagents generated in this study are available upon request from the Lead Contact with a completed Materials Transfer Agreement.

#### Data and Code Availability

Code used for statistical analysis and RNA-seq, ATAC-seq, MeDIP-Seq data generated in this study will be publicly available as of the date of publication. Raw sequencing data will be available on GEO. Original data for all Western blots, gels, and other figures will be available from the Lead Contact upon request. This paper does not report original code. Software packages used are listed in the Key Resources Table and described in the Method Details section.

### EXPERIMENTAL MODEL AND SUBJECT DETAILS

#### Overview of mouse cohorts and experimental design

We used middle-aged (15-month-old) C57BL/6JN mice obtained from the National Institute on Aging (NIA) for all *in vivo* studies. Animals were maintained under a 12-hour light/dark cycle with ad libitum access to food and water. Upon arrival, the mice were housed in groups of 3–4 per cage and closely monitored for well-being. Following an acclimatization period, animals were randomly and blindly assigned to different treatment groups. All experimental procedures involving mice were approved by the University Animal Research Committee (UCAR) at the University of Rochester and were performed in compliance with institutional guidelines.

#### Aging study cohort and fucoidan administration

The aging study comprised 80 mice (40 male, 40 female). 44 mice received fucoidan in their diet, and the remaining 36 were fed a control diet. *F. vesiculosus* fucoidan for *in vivo* mouse studies was obtained from DoNotAge.org. Fucoidan was administered to the mice using two methods – by adding it to their drinking water and by mixing it into their chow. For the drinking water, we prepared Fucoidan at a concentration of 10 g/L. The drinking water was changed three times a week. Since each mouse typically drinks 5 mL of water per day, they received roughly 50 mg of fucoidan daily from this route. At the same time, fucoidan was mixed into the diet by combining 8 g of the compound with 150 g of dry chow. With adult mice eating around 4-5 g of chow per day, this results in an intake of approximately 203-253 mg of fucoidan per day. Overall, the mice received a total daily dose of about 253-303 mg of fucoidan per day, averaging around 278 mg of fucoidan per day. Frailty was scored once a month beginning at 19 months of age and continuing until they were 25 months old. Frailty scoring was performed in a blinded manner, as described in by Whitehead *et al*^33^. Blood samples were collected at 22 and 26 months of age.

#### Tissue collection cohort

Another 20 mice (5 males and 5 females on fucoidan diet, and 5 males and 5 females on control diet) were euthanized after one month on their respective diets. Tissues were harvested and utilized for RNA-seq, ATAC-Seq, MeDIP-Seq, and protein analysis by Western blotting.

#### Paraquat challenge cohort

The cohort used for paraquat challenge comprised 24 mice (6 per diet treatment group per sex). After receiving fucoidan or control diets for one month, these mice were injected intraperitoneally with either PBS (control) or paraquat at a dose of 50 mg/kg (LD_75/15_ and were subsequently euthanized. Tissues collected from these animals were used for RT-qPCR.

#### Sirt6 KO cohort

SIRT6 KO mice were obtained from JAX, Strain #:006050. These mice are maintained in Rochester as heterozygous animals and then bred to homozygosity for the experiments. Breeding pairs were randomly assigned to one of two dietary regimens: a control group receiving standard chow and water or a treatment group receiving chow and water supplementation with fucoidan at an average daily dose of approximately 278 mg/mouse/day. Following mating, mothers maintained their respective diet throughout gestation. Upon parturition, neonatal pups were administered up to 100 uL of the corresponding water (control or fucoidan-supplemented) daily via a flexible 1 mL pipette while mothers continued their assigned diet. The survival of the Sirt6 KO pups was monitored closely.

### METHOD DETAILS

#### Chemicals

Fucoidan was obtained from the following sources. Sigma-Aldrich: Fucoidan from *F. vesiculosus* (F8190 and F5631), Fucoidan from *U. pinnatifida* (F8315), Fucoidan from *M. pyrifera* (F8065), Fucoidan from DoNotAge.org. and Kanahide Bio Okinawa Fucoidan (4958349250135). L-Fucose was obtained from Sigma-Aldrich (F2252), MDL-800 from Cayman Chemical Company (35528) and MDL-801 from Aobious (AOB17342).

#### Purification of recombinant SIRT6

Chemically competent Rosetta Gami B (DE3) (Millipore) *E. coli* were transformed with a pET11a plasmid encoding N-terminally 6×His-tagged SIRT6. Bacteria were grown at 37°C in LB medium supplemented with 100 µg/mL ampicillin, 25 µg/mL kanamycin, and 15 µg/mL chloramphenicol, reaching an OD_600_ of approximately 2.0. Protein production was induced with 0.4 mM IPTG, after which the culture was shifted to room temperature (RT) and incubated for 20 h. Cells were then pelleted by centrifugation, flash-frozen in liquid nitrogen, and stored at –80°C.

Frozen bacterial pellets were thawed on ice and resuspended in a lysis buffer composed of 50 mM Tris (pH 7.5), 500 mM NaCl, 10 mM imidazole, 10 mg lysozyme, 0.1% Triton X-100, and one Roche complete protease inhibitor tablet. The cell suspension was sonicated briefly, rotated at 4°C for 1 h, and then sonicated again before clarification by centrifugation at 15,000 × *g* for 1 hour at 4°C. The supernatant was incubated with Ni-NTA agarose beads (Qiagen) that had been equilibrated with wash buffer (50 mM Tris, pH 7.5; 500 mM NaCl) for 1 hour at 4°C. Beads were pelleted by centrifugation and washed with 50 column volumes of wash buffer containing 50 mM imidazole. SIRT6 was eluted with the same buffer supplemented with 0.5 M imidazole for 15 min at 4°C on a rotator. The eluted protein was filtered, dialyzed overnight at 4°C in 10 kDa MWCO SnakeSkin dialysis tubing (Thermo) against 50 mM Tris (pH 7.5), 150 mM NaCl, 5% glycerol, and 1 µM ZnCl_2_, then filtered again, flash-frozen as drops in liquid nitrogen, and stored at –80°C.

#### SIRT6 self-ribosylation assay

BL21 (DE3) *E. coli* were transformed with a plasmid encoding 6×His-tagged NMNAT1 (NMNAT1_OHu18148C_pET24a, Genscript). A single colony was grown overnight in 50 mL LB at 37°C, after which 10 mL of this culture was used to inoculate 1 L of LB in baffled flasks. Cells were grown at 37°C to an OD600 of approximately 0.7– 1.0, induced with 2 mM IPTG, and incubated for an additional 3 h before pelleting and flash-freezing at –80°C. Thawed cells were resuspended in 50 mM NaPO_4_ (pH 7.4), 300 mM KCl, 1 mM MgCl_2_, 1 mM DTT, 10 mg lysozyme, 0.1% Triton X-100, 1 Roche protease inhibitor tablet, and 10 mM imidazole, and rotated at 4°C for 1 h. The lysate was sonicated and clarified by centrifugation, and the supernatant was incubated with Ni-NTA beads at 4°C for 1 h. Following washes with buffer containing 50 mM imidazole, NMNAT1 was eluted with buffer containing 500 mM imidazole. The eluted protein was filtered to 0.45 µm and dialyzed overnight at 4°C in 25 mM NaPO_4_ (pH 7.4), 150 mM KCl, 500 µM MgCl_2_, 500 µM DTT, and 50% glycerol, then stored at –20°C.

P-NAD+ was synthesized by mixing 500 ng of purified NMNAT1 with 1.665 µM ATP (α-^32^P), 1.665 µM β-NMN, and NMNAT1 buffer (50 mM KPO_4_, pH 7.4; 300 mM KCl; 30 mM MgCl_2_; 1 mM DTT) at 37°C for 1 h. The reaction was subsequently stored at – 20°C.

SIRT6 self-ribosylation reactions contained 3 µg of purified SIRT6, 100 µM non-radioactive NAD+ supplemented with 100 nM ^32^P-NAD+, 1 mg/mL fucoidan product (activator), and SIRT6 buffer (50 mM Tris, pH 7.5; 150 mM NaCl). A master-mix without the activator was prepared on ice, then aliquoted into PCR tubes containing the fucoidan. Samples were mixed and incubated at 37°C for 1 h. Reactions were quenched by boiling for 5 min in 1× Laemmli buffer containing 5% β-mercaptoethanol. Proteins were separated by SDS-PAGE (precast gels, Bio-Rad) at 240 V for 30 min in 1× TGS buffer and transferred onto PVDF membranes (midi format) using a Trans-Blot Turbo system (Bio-Rad). Membranes were washed (3 × 5 min in 1× TBST), air-dried, and exposed to a phosphor storage screen. Imaging was performed on an Azure Sapphire phosphor imager, and band intensities were quantified in ImageJ. A duplicate gel was stained with Coomassie to confirm equal loading.

#### Mononucleosome deacetylation assay

Mononucleosomes (2 µg H3K9Ac; Epicypher) were incubated with 2 µg purified SIRT6, 1 mg/mL fucoidan product, and 500 µM NAD+ in a total volume sufficient for 2 h at 37°C. The reaction was quenched by boiling in 1× Laemmli buffer containing 5% β-mercaptoethanol for 5 min. The samples were split onto two precast SDS-PAGE gels (Bio-Rad). One gel was stained with Coomassie for loading control, and the other was transferred to nitrocellulose for Western blotting. The membrane was blocked in 5% BSA in TBST for 1 h at room temperature, washed, and probed with mouse anti-H3K9Ac antibody (ab4441, Abcam) before incubation with goat anti-mouse HRP (ab6721, Abcam). ECL reagents (Bio-Rad) were used to visualize bands on a Bio-Rad ChemiDoc system, and signal intensities were quantified with Bio-Rad Image Lab.

#### Blood and plasma collection

Blood samples were centrifuged at 2000 × *g* for 15 min at 4°C to separate plasma, which was used for cytokine profiling. The remaining cell pellet was used for DNA and RNA extraction for measuring DNA methylation age and for total RNA sequencing.

#### Plasma cytokine profiling

The plasma samples were sent to Luminex Corp. for chemokine and cytokine profiling using the ProcartaPlex Mouse Cytokine and Chemokine Panel 1A, 36-Plex (Thermo Fisher, Catalog #EPX360-26092-901) and the associated Control Set (Catalog #EPX360-26092-CTR).

#### DNA and RNA isolation from blood

Pelleted blood cells were mixed with 1× Monarch DNA/RNA Protection Reagent (NEB, T2011L) and processed using the NEB Total RNA Extraction Kit (T2010) according to the manufacturer’s protocol. The genomic DNA was eluted from the gDNA removal column, and total RNA was collected from the RNA purification column.

#### Epigenetic Clock Analysis

DNA samples from blood were sent to the Clock Foundation for epigenetic age prediction using the UniversalClock3Blood.

#### RNA isolation from frozen tissues

Tissues were pulverized to a fine powder before RNA extraction. All samples and materials were kept on liquid nitrogen to prevent RNA degradation. RNA was isolated from pulverized tissue using Trizol/Chloroform extraction. 500 µl – 1 mL Trizol (Invitrogen, 15596018) was added to each sample, vortexed, and incubated at room temperature for 5 minutes. 100-200 µl chloroform was added to the supernatant, shaken vigorously for 15 seconds, incubated at room temperature for 3-5 minutes, then centrifuged at 12,000 x g for 15 minutes. Aqueous phase containing RNA was carefully separated from the rest of the sample and mixed 1:1 with isopropanol before incubating at room temperature for 10 minutes and then spinning at 12,000 x g for 10 minutes. RNA pellet was then washed twice with 75% ethanol in nuclease-free H2O and centrifuged at 8,000 x g for 5 minutes. Pellet was allowed to air dry before resuspending in nuclease-free H2O.

#### RT-qPCR

RNA was further purified using RNA PureLink Mini Kit (Thermo, 12183020). cDNA was generated from RNA using iScript cDNA Synthesis Kit (BioRad, 1708891). qRT-PCR was performed using BioRad CFX Connect Real Time machine and SYBR Green Master Mix (BioRad, 1725150) using 30ng cDNA per reaction. Each sample was run in triplicates. Mouse L1 primers were targeted specifically towards L1MdA elements as previously described. Primers used as follows: mL1 fwd - ATGGCGAAAGGCAAACGTAAG; mL1 rev - ATTTTCGGTTGTGTTGGGGTG; mEEF2 fwd - GATGATCACCATCCACTTACC; mEEF2 rev - GGGTCGCAGCTCTTAATAC; p21 fwd – CGAGAACGGTGGAACTTTGAC; p21 rev – CAGGGCTCAGGTAGACCTTG.

#### RNA-seq library preparation

RNA sequencing libraries were prepared from total RNA using the TruSeq Stranded Total RNA Library Prep Kit (Illumina, 20020697) by the University of Rochester Genomics Research Center. Sequencing was performed on an Illumina NovaSeq platform, producing 150 bp paired-end reads with a sequencing depth of 60 million reads per sample.

#### ATAC-Seq library preparation

ATAC sequencing was performed similaily to the basic ATAC-sequencing protocol^46^. Briefly, frozen tissues were processed using the Chromium Nuclei Isolation Kit (10X Genomics, 1000493). 50,000 isolated nuclei were pelleted and resuspended in 1× TD buffer, 16.5 µL 1× PBS, 0.5 µL 10% Tween-20, 0.5 µL 1% Digitonin, 2.5 µL Tn5 transposase (Diagenode, C01070012), and nuclease-free H_2_O to a total volume of 25 µL. The mixture was incubated at 37°C for 30 min with shaking at 1000 rpm. Transposed DNA fragments were purified using the MinElute PCR Purification Kit (Qiagen, 28006). Libraries were amplified using NEBNext High Fidelity 2× PCR Master Mix (NEB, M0541L) and pre-mixed primers bearing unique dual indexes (IDT). A double-sided SPRI bead cleanup (SPRIselect, Beckman Coulter, B23318) was employed for final size selection of the libraries. Sequencing was conducted at the University of Rochester Genomics Research Center using an Illumina NovaSeq platform, producing 75 bp paired-end reads with a sequencing depth of 50 million reads per sample.

#### MeDIP-seq library preparation

Genomic DNA samples were extracted from tissues using the DNeasy Blood & Tissue Kit (Qiagen, 69506), according to the manufacturer’s instructions. 1 ug extracted genomic DNA was randomly sheared to a median fragment size of 200 base pairs by sonication using a Bioruptor, set at 30% amplitude for 12 cycles, with each cycle consisting of 30 seconds ON and 30 seconds OFF. The fragmented DNA was then processed using the NEBNext Ultra II DNA Library Prep Kit for Illumina (New England Biolabs, E7600S), which involved end repair, 3′ adenylation to create a 3′ dA overhang, and adapter ligation. Size selection of adapter-ligated DNA was performed with Ampure XP beads, targeting fragments ranging from 200 to 250 base pairs. Methylated DNA was subsequently immunoprecipitated using the Zymo Methylated-DNA IP Kit (D5101), employing a 1:10 ratio of DNA to antibody to ensure specificity. The library was PCR amplified for 15 cycles. Sequencing was conducted at the University of Rochester Genomics Research Center using an Illumina NovaSeq platform, producing 150 bp paired-end reads with a sequencing depth of 50 million reads per sample.

#### Western blot

Tissues were pulverized using a liquid nitrogen cooled tissue crusher. Cells were lysed with a 2x Laemmli solution and incubated 10 min at 90℃, followed by centrifugation at 13,000g for 10min to remove debris, and the supernatant was transferred to a new tube. Samples were loaded into a BioRad Criterion 4–20% gel. After transfer to PDVF membrane and blocking (5% BSA) for 2 h at RT, membranes were incubated overnight with primary antibodies in 5% blocking buffer at 4°C. Membranes were washed 3x with TBST for 5 min each before the secondary antibody in 1x TBST was added for 1h incubation at RT. Membranes were washed 3x with TBST for 5 min and then imaged. The following antibodies were used: H3 (CST, 12648S)-1:5,000, H3K9ac (Abcam, ab4441)-1:1,000, and H3K9me3 (Abcam, ab8898)-1;1000.

### QUANTIFICATION AND STATISTICAL ANALYSIS

#### Gel quantification

Radioactive signals (phosphorimaging) on blots were quantified with ImageJ. Western blot signals were quantified using Bio-Rad Image Lab, with Coomassie-stained gels serving as loading controls.

#### Statistics

Statistical analysis as performed in R or GraphPad Prism. Statistical tests are specified in the text and figure legends. The overall significance of frailty differences between fucoidan and control treated mice was determined with a linear mixed model with formula Frailty∼Month*Diet+(1|AnimalID), where Months denotes the time since the beginning of treatment. Significance stars use the following convention:. p < 0.1, * p < 0.05, ** p < 0.01, *** p < 0.001. Multiple testing corrections is performed using the Benjamini-Hochberg procedure where specified.

#### RNA-seq analysis

Reads were trimmed using TrimGalore! and aligned to the mouse genome GRCm39 using STAR^47^, with parameters --outFilterMultimapNmax 100 -- winAnchorMultimapNmax 200 --outFilterMismatchNoverLmax 0.04. Gene and transposon expression was simultaneously quantified using TEtranscripts^48^, with gene annotations sourced from Ensembl build 108 and TE annotations from the TEtranscripts website. Only transcripts with the “protein_coding” biotype were included for analysis of the aging cohort blood data. Differential expression was performed using EdgeR^49^ after removing low expression genes and TEs using filterByExpr. GSEA^50^ and ORA were performed using ClusterProfiler^51^. Cell population deconvolution was performed using mMCP-counter^34^ made available by the immunedeconv R package^52^. Aging coefficients for tabula muris data were computed using EdgeR based the count tables provided by the authors^39^, including animals 12 mo or older to restrict to ages similar to those of the animals in our study. Differential expression for Sirt6 OE vs control was also based on the count tables provided by the authors^8^.

#### ATAC-seq analysis

Read were trimmed using TrimGalore! and aligned to the mouse genome GRCm39 using Bowtie2^53^ with parameters --very-sensitive -X 1000 –dovetail. Improper pairs and secondary alignments were discarded using samtools^54^ view -f 0×2 -F 0×100. This has the effect of retaining only one random alignment among multimapping reads, as recommended by Teissandier et al.^55^ Mitochondrial reads and PCR duplicates were removed with samtools and picard. Cleaned bam files were converted to bed format and used to call peaks with MACS2^56^ with parameters -f BED --keep-dup “all” -q 0.01 -- nomodel --shift −100 --extsize 200 for each sample individually. A unified peak set was generated by first merging the peaksets of all samples using bedtools^57^ merge. This step was performed separately for lung and liver samples. Next, we identified regions which were called as peak in at least 2 samples using bedtools multiinter. Finally, we only kept peaks within the union peakset if they contained at least one of the regions called in 2 or more samples of the same tissue using bedtools intersect. Repetitive element annotation was performed with RepeatMasker. Accessibility of OCRs and repetitive elements was simultaneously quantified using featureCounts in the subread package, with parameters -F SAF -p -B --read2pos 5 -O --fraction, which reduces reads to their 5’ end (Tn5 cut site) and counts reads fractionally, if they overlap both a peak region and a repeat region. Differential accessibility was evaluated using EdgeR^49^.

#### MeDIP-Seq analysis

Reads were trimmed with Fastp^58^ to remove adapters, low-quality bases, and Illumina-specific sequences. High-quality reads were aligned to the mouse genome (GRCm38) using STAR^47^ with parameters –outFilterMismatchNmax 3 and -- outFilterMultimapNmax 100 to account for multiple mappings of TEs. TEtranscripts^48^ was used to quantify MeDIP read counts over annotated TE regions available on the TEtranscript website. DESeq2^59^ was then employed to identify TE families that were hyper- or hypomethylated in fucoidan-treated mice compared to controls.

#### Overall trend of LINE1 expression / epigenetic state

Following differential analysis in RNA-seq, ATAC-seq or MeDIP-seq, a binomial test was applied to evaluate whether the distribution of up or downregulation events deviated significantly from 50%. TE families with |log_2_FoldChange)| > 0.25 and *p* < 0.05 (unadjusted) were highlighted in volcano plots.

## Author Contributions

EH purified recombinant proteins and conducted *in vitro* assays. AB coordinated animal feeding, lifespan analysis, frailty analysis and blood collections. VP and Z. Zhang assisted in lifespan experiments and blood collections. VP performed frailty scoring in a blinded manner. CR isolated RNA and DNA from blood of aging cohort. FM did statistical analyses on lifespan, frailty, epigenetic clock and cytokine data. CR isolated tissues, isolated RNA and prepared ATAC-seq libraries from 1-month-treated animals. ML did Western blotting on tissues and prepared MeDip-seq libraries. FM performed data processing for RNA-seq and ATAC-seq and analyzed RNA-seq and cytokine abundance. VV validated RNA-seq analyses. Z. Zheng performed data processing for MeDip-seq and analyzed transposable element expression, accessibility and methylation status. YL performed paraquat treatment and isolated tissues from paraquat-treated animals. JM and NS performed RT-PCR experiments. VG, AS, FM, CR and AB assembled the manuscript with input from all authors. GT, DF and MS provided scientific input. VG and AS supervised the study.

## Acknowledgements

This research is supported by grants from US National Institute on Aging to AS and VG.

---

**Figure S1: Body mass and frailty score breakdown by major system.**

**(A)** Body mass of mice fed Fucoidan or control diets over time. **(B)** Breakdown of frailty scores by major organ / system, and number of mice per group.

**Figure S2: Evidence of Sirt6 activation mediating the beneficial effects of fucoidan.**

**(A)** Lifespan curve for Sirt6 KO mice fed with fucoidan or control diets. **(B)** Western blot against the Sirt6 substrate H3K9ac (3-4 biological replicates per group). P values were calculated using t test.

**Figure S3: RNA-seq analysis of blood in female mice.**

**(A)** Volcano plot for the effect of fucoidan on 22-month female mice, which were part of the survival study. **(B)** GSEA results showing GO terms significantly enriched in female mice. **(C)** Plasma levels of cytokines in female mice measured by ELISA. **(D)** Estimation of blood cell composition in female mice by deconvolution using mMCP-counter. p-values were calculated with a Mann-Whitney U test.

**Figure S4: Effects of fucoidan treatment common across tissues and sexes.**

**(A)** Upset plot showing intersections of sets of genes upregulated for each sex and tissue. Empty intersections not shown. Selected intersections were tested for enrichment of Gene Ontology biological processes. Intersections were denoted by the same color in the upset plot and enrichment results. **(B)** Upset plot showing intersections of sets of genes downregulated for each sex and tissue. Empty intersections not shown. Selected intersections were tested for enrichment of Gene Ontology biological processes. Intersections were denoted by the same color in the upset plot and enrichment results.

**Figure S5: Additional analysis of gene and transposon expression in lung and liver tissues.**

**(A, B, C)** Comparison of the effects of fucoidan treatment vs the effect of aging in the tissue and sex specified. Darker dots represent genes differentially expressed in both Fucoidan vs control and during aging (adjusted p-value < 0.05 for aging, adjusted p-value < 0.1 for Fucoidan vs control). **(D)** Comparison of the effects of fucoidan treatment and Sirt6 overexpression in female liver tissue. Darker dots represent genes differentially expressed in both Fucoidan vs control and Sirt6 OE vs control (adjusted p-value < 0.05, |logFC| > 1 for both). **(E)** RNA-seq, ATAC-seq and MeDIP-seq volcano plots for the effect of fucoidan treatment on LINE1 element expression, accessibility and methylation in liver. TE families with logFC < −0.25 and p < 0.05 (unadjusted) are colored in blue. TE families with logFC > 0.25 and p < 0.05 (unadjusted) are colored in red. Overall asymmetry of the volcano plot is tested for significant deviations from equal likelihood (50%) using a binomial test.

