## Supplementary Figures for "SIRT6 activator fucoidan extends healthspan and lifespan in aged wild-type mice"

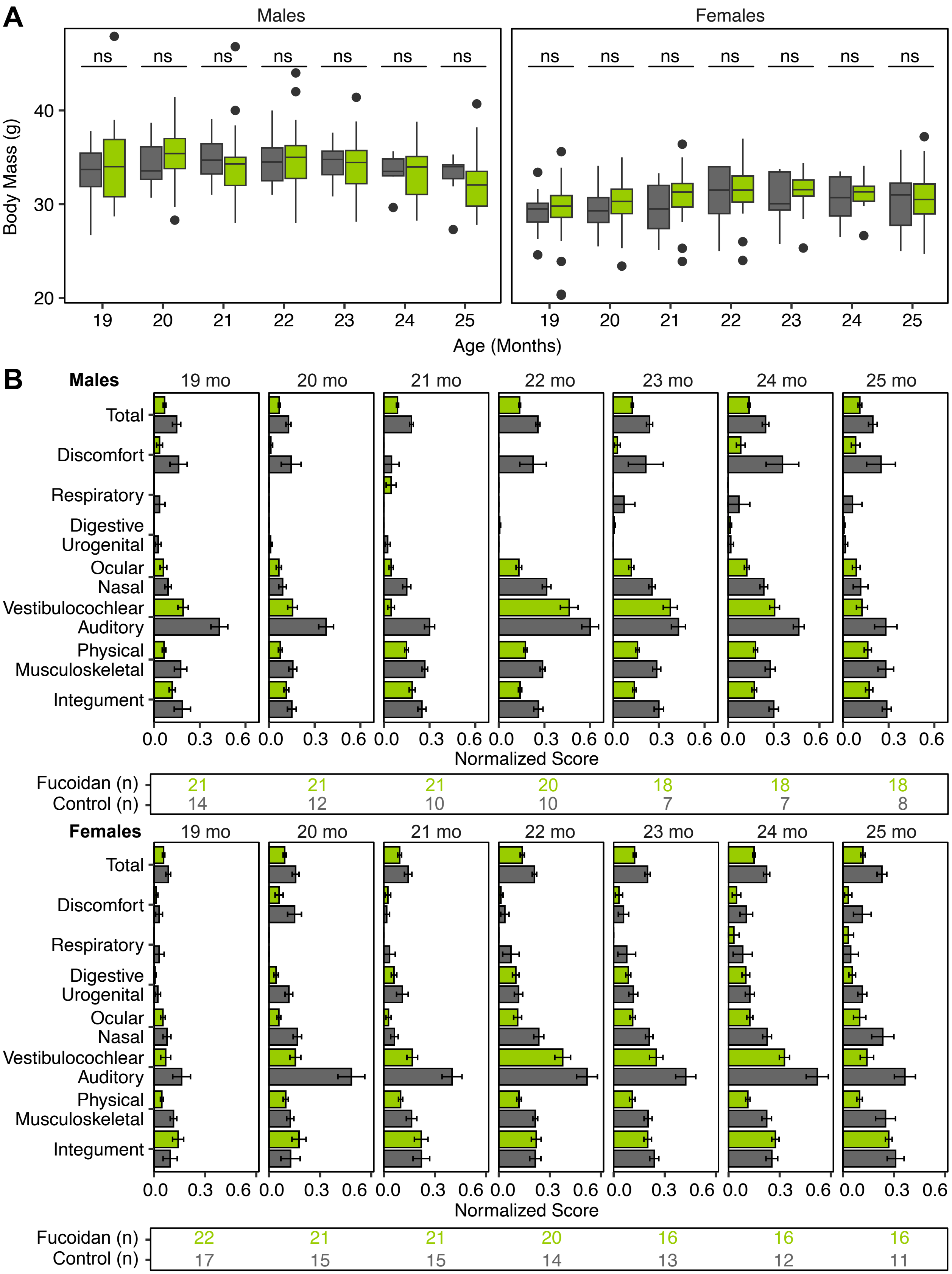


**Figure S1: Body mass and additional frailty score information. (A)** Body mass of mice fed Fucoidan or control diets over time. **(B)** Breakdown of frailty scores by major organ / system, and number of mice per group**.**

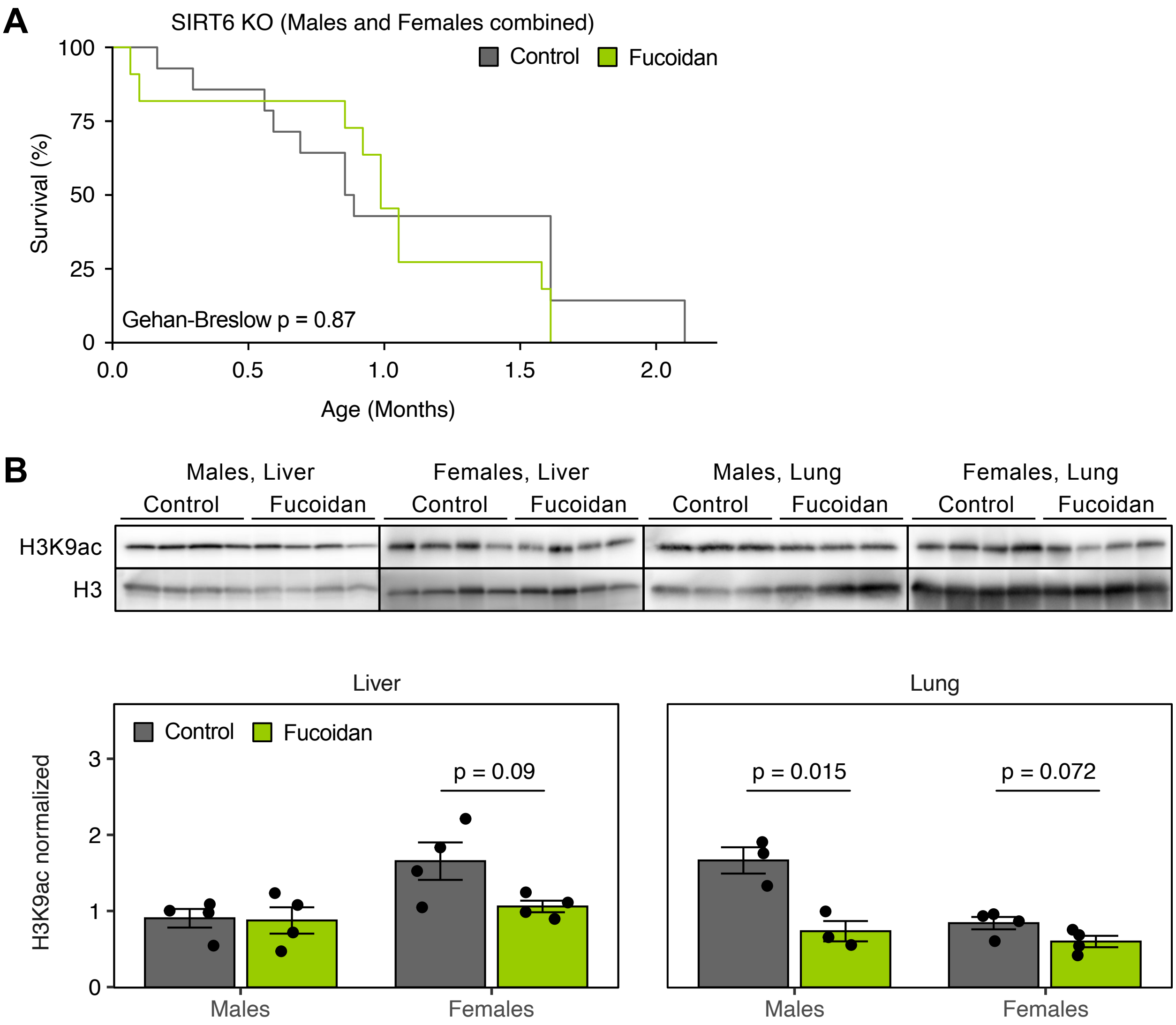


**Figure S2: Evidence of Sirt6 activation mediating the beneficial effects of fucoidan. (A)** Lifespan curve for Sirt6 KO mice fed with fucoidan or control diets. **(B)** Western blot against the Sirt6 substrate H3K9ac.

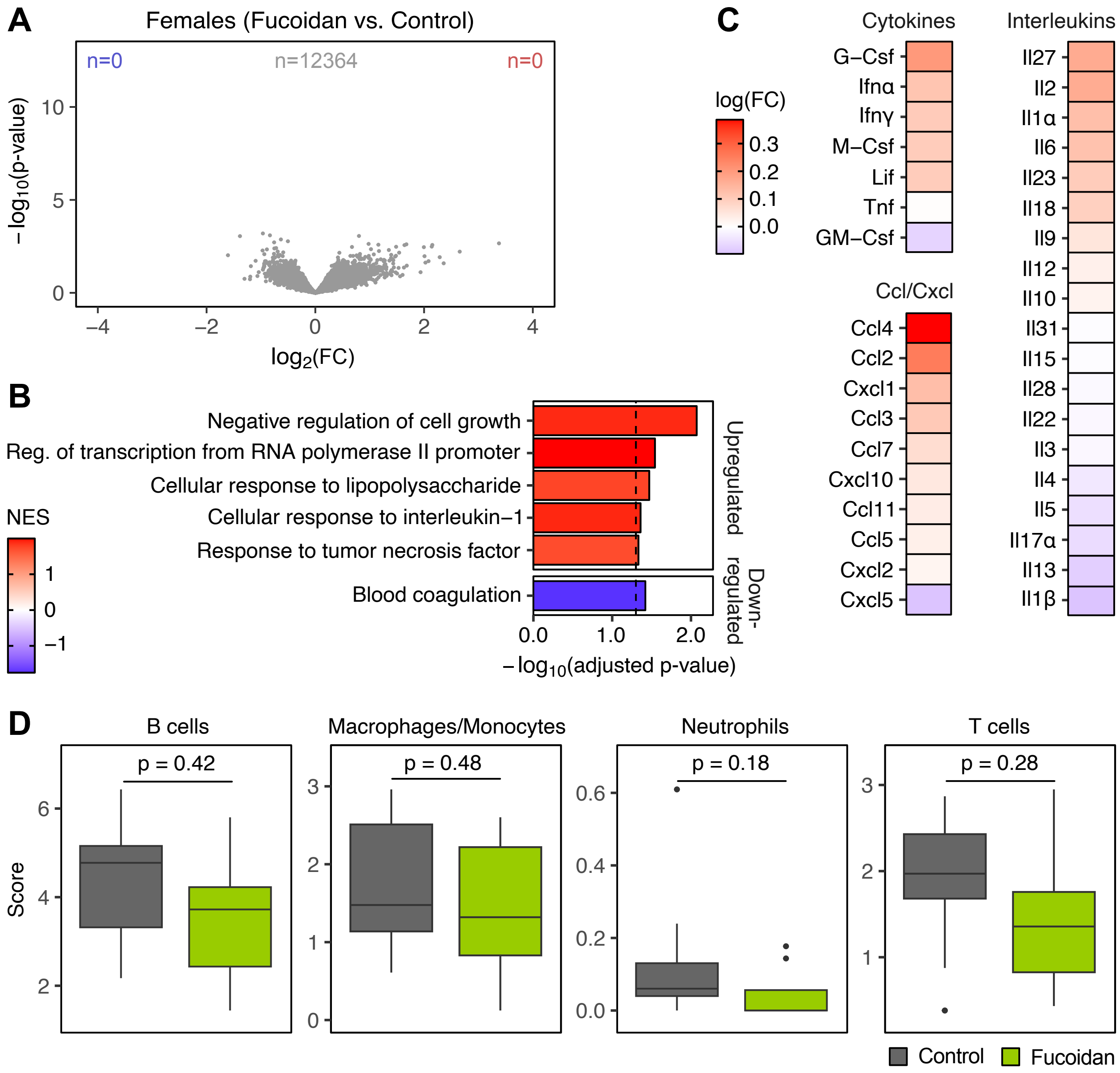


**Figure S3: RNA-seq analysis of blood in female mice. (A)** Volcano plot for the effect of fucoidan on 22 mo female mice, which were part of the survival study. **(B)** GSEA results showing GO terms significantly enriched in female mice. **(C)** Plasma levels of cytokines in female mice measured by ELISA. **(D)** Estimation of blood cell composition in female mice by deconvolution using mMCP-counter. P-values were calculated with a Mann-Whitney U test.

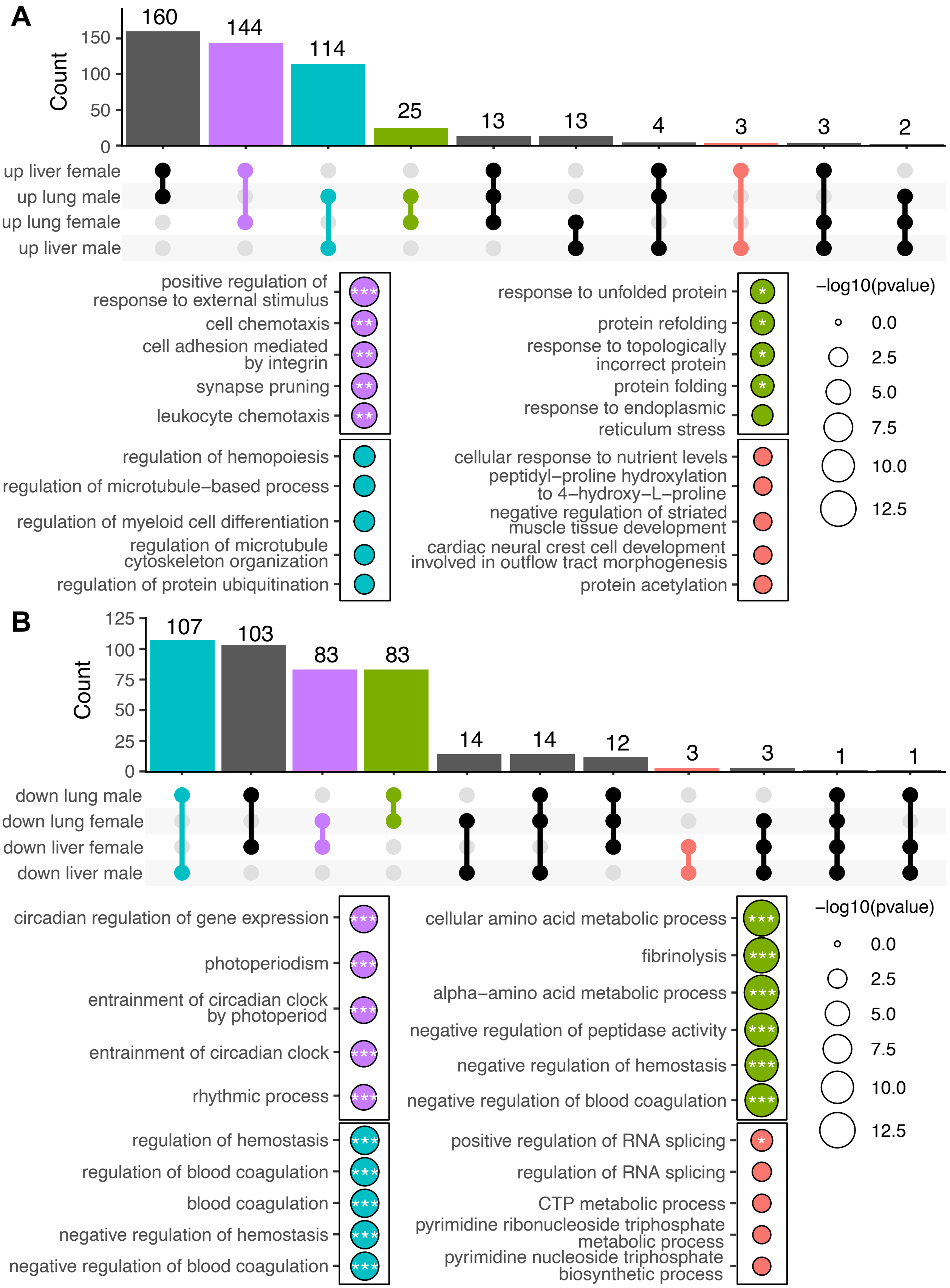


**Figure S4: Effects of fucoidan treatment common across tissues and sexes. (A)** Upset plot showing intersections of sets of genes upregulated for each sex and tissue. Empty intersections not shown. Selected intersections were tested for enrichment of Gene Ontology biological processes. Intersections were denoted by the same color in the upset plot and enrichment results. **(B)** Upset plot showing intersections of sets of genes downregulated for each sex and tissue. Empty intersections not shown. Selected intersections were tested for enrichment of Gene Ontology biological processes. Intersections were denoted by the same color in the upset plot and enrichment results.

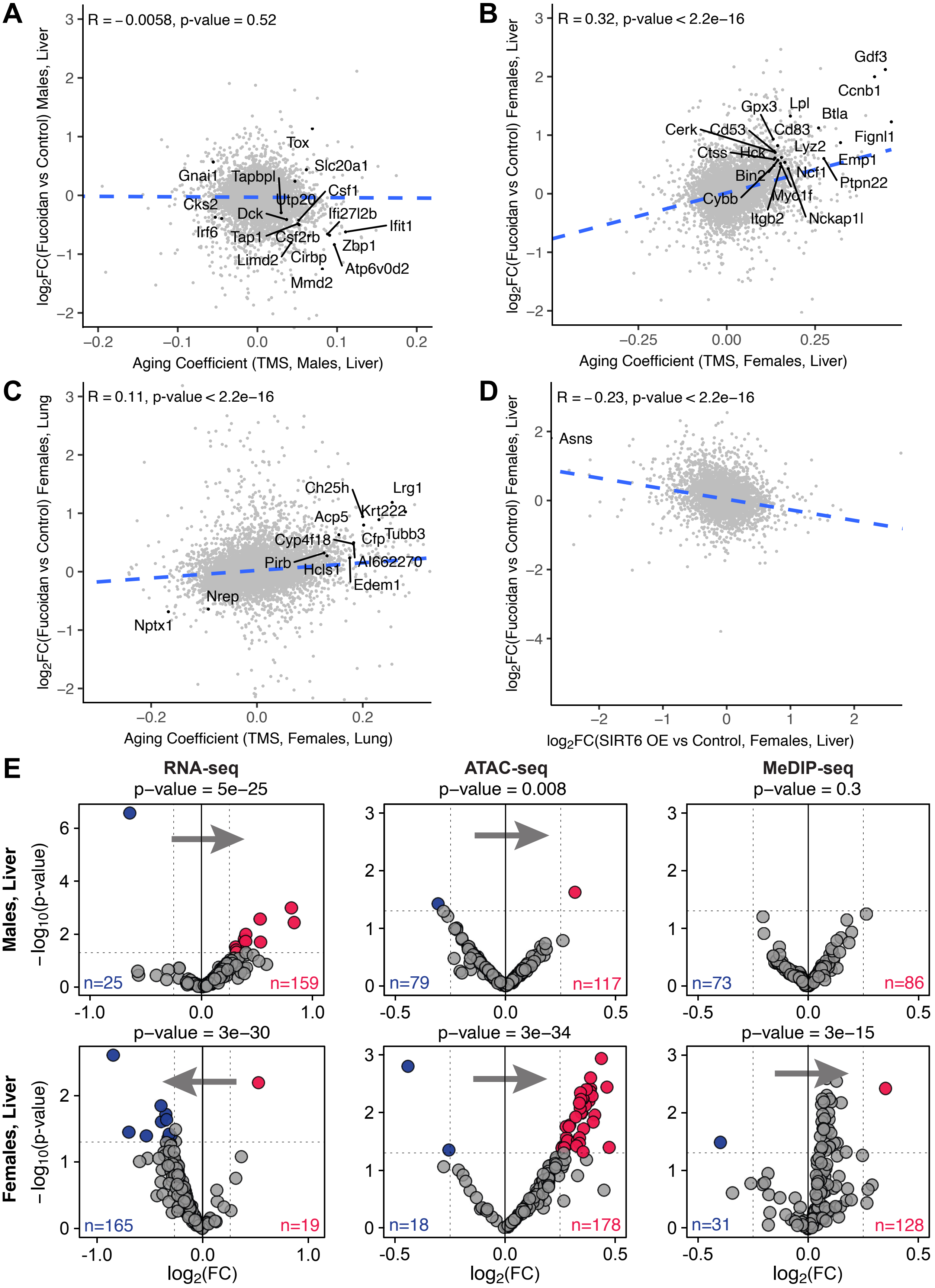


**Figure S5: Additional analysis of gene and transposon expression in lung and liver tissues. (A, B, C)** Comparison of the effects of fucoidan treatment vs the effect of aging in the tissue and sex specified. Darker dots represent genes differentially expressed in both Fucoidan vs control and during aging (adjusted p-value < 0.05 for aging, adjusted p-value < 0.1 for Fucoidan vs control). **(D)** Comparison of the effects of fucoidan treatment and Sirt6 overexpression in female liver tissue. Darker dots represent genes differentially expressed in both Fucoidan vs control and Sirt6 OE vs control (adjusted p-value < 0.05, |logFC| > 1 for both). **(E)** RNA-seq, ATAC-seq and MeDIP-seq volcano plots for the effect of fucoidan treatment on LINE1 element expression, accessibility and methylation in liver. TE families with logFC < -0.25 and p < 0.05 (unadjusted) are colored in blue. TE families with logFC > 0.25 and p < 0.05 (unadjusted) are colored in red. Overall asymmetry of the volcano plot is tested for significant deviations from equal likelihood (50%) using a binomial test.
